# Derepression of transposable elements in mouse prefrontal cortex disrupts social behavior

**DOI:** 10.1101/2025.03.31.646358

**Authors:** R. Kijoon Kim, Corinne Smith, Natalie L. Truby, Nic Carwile, Gabriella M. Silva, Rachael L. Neve, Xiaohong Cui, Peter J. Hamilton

## Abstract

Here, we present a synthetic biology approach to assess the social behavioral consequences of altered function of the Krüppel-associated box zinc finger protein (KZFP) interacting protein TRIM28 within the prefrontal cortex (PFC) of male and female mice. We reprogrammed natural TRIM28^WT^ by replacing the transcriptionally repressive domain with an enhanced transcriptional activation domain VP64-p65-Rta (TRIM28^VPR^), or by excising the transcriptional regulatory domain (TRIM28^NFD^). *In vitro* validation confirmed that TRIM28^WT^ represses, and TRIM28^VPR^ activates, the expression of a KZFP-regulated *luciferase* reporter gene. Upon intra-PFC viral-mediated delivery of TRIM28 variants, we observed that inversion of TRIM28 transcriptional control via HSV-TRIM28^VPR^ reduced the salience of novel social interaction for male and female mice while not affecting non-social behaviors. RNA-sequencing revealed HSV-TRIM28^VPR^ promoted transcriptional escape of all classes of TEs, particularly those located within intronic and distal enhancer regions of downregulated immune genes. HSV-TRIM28^VPR^-driven social deficits were reversible by intra-PFC repletion of interferon cytokines. These novel data point to PFC KZFP-TRIM28 interactions as necessary to stabilize TEs to enable cis-regulation of key immune gene expression and enhance organismal capacity for complex, pro-social behaviors.

## Introduction

Social behaviors are a fundamental aspect of group living and play substantial roles in determining the health and wellbeing of both the individual and society^1–3^. Dysfunction in social behaviors is a hallmark of nearly all neuropsychiatric syndromes. For example, social withdrawal characterizes conditions including major depressive disorder, autism spectrum disorder (ASD), and schizophrenia, while hyper-sociability occurs in Angelman and Williams syndromes^4^. However, the neurobiological mechanisms that enable complex social behaviors are incompletely understood.

Since their discovery by Barbara McClintock, genomic transposable elements (TEs), or self-replicating DNA sequences, have been identified as molecular drivers capable of altering genome size and stability and are thought to contribute to the diversification and evolution of species^5–8^. However, due to the potential deleterious effects of active transposition of TEs, maintaining a transcriptionally silenced TE landscape is vital to the health of host organisms^9–13^. Genomic regions consisting of transcriptionally silenced TEs have been co-opted by their hosts in diverse ways, including in formation of cis-regulatory regions^14–16^ and by contributing to the regulation of the adaptive immune system^17–19^. Recently, several groups have begun to identify evidence that links social and other complex behaviors with proper regulation of these TE-linked immune responses in the brain^20–22^.

Krüppel-associated box (KRAB) zinc finger proteins (KZFP) comprise an evolutionarily ancient and highly diverse family of transcription factors (TF) that have evolved to limit TE transposition by repressing TE expression via the recruitment of tripartite motif protein 28 (TRIM28), also called KAP1^23–27^. KZFP-conserved KRAB domains recruit TRIM28 through its RBCC motif, and TRIM28 recruits histone methyltransferases (HMTs) including SETDB1 and the nucleosome remodeling and deacetylase (NuRD) complex via the TRIM28 HP1, PHD, and BROMO domains^28,29^. This mechanism contributes to the formation of heterochromatin at KZFP-DNA binding sites and transcriptional silencing of KZFP gene targets, including TEs^30^.

Here, we utilized a synthetic biology approach to analyze the transcriptional and behavioral consequences of TRIM28-mediated transcriptional activity and dysfunction in the brain. In addition to the endogenous repressive TRIM28^WT^ protein, we created novel synthetic TRIM28 variants with intact KZFP-binding domains, each designed to exert distinct forms of transcriptional control across KZFP-regulated genomic loci. First, we replaced the repressive HP1, PHD, and BROMO domains with the transcriptional activator VP64-p65-Rta (VPR) to create the novel synthetic TRIM28^VPR^. We also excised these repressive domains to create the transcriptionally inert TRIM28^NFD^. Following packaging into herpes simplex viruses (HSVs), we delivered our HSV-TRIM28 variants versus HSV-GFP control virus to male and female mouse prefrontal cortex (PFC) and conducted a battery of behavioral and transcriptomic assays to determine the behavioral and molecular consequences of dysregulating TRIM28 transcriptional control in PFC.

In both male and female test mice, we observed that intra-PFC delivery of HSV-TRIM28^VPR^ induced significant alterations to social behaviors, characterized by a lack of interest in novel social interactions and impaired ability to engage in previously established social hierarchies, without producing detectable non-social behavioral changes. In RNA-sequencing (RNAseq) analyses, HSV-TRIM28^VPR^ activated TE escape and downregulated immune-related genes located in genomic regions associated with these released TEs. Ablation of TRIM28-mediated transcription through delivery of HSV-TRIM28^NFD^ induced similar, but modest social behavioral deficits and transcriptional changes, pointing to a graded relationship between magnitude of PFC TE dysregulation and degree of social deficits. These social deficits could be restored by introduction of exogenous immune factors to manipulated PFC. In sum, this body of work indicates that TRIM28 participates in the transcriptional stabilization of TE-rich genomic regions, and dysregulation of this TRIM28-mediated transcriptional control in PFC allows TE escape, impedes the expression of primarily immune-related genes, and diminishes the capacity of the organism to perform complex social behaviors via disruption of PFC immune gene expression. This points to a novel mechanistic relationship between cortical TEs, immune functions, and social cognition.

## Results

### Synthetic TRIM28 variants bi-directionally control the expression of a KZFP-regulated gene *in vitro*

Here, we employed a synthetic biology approach to investigate the gene-regulatory functions of TRIM28. We cloned three different synthetic TRIM28 variants into overexpression plasmids: TRIM28^WT^, identical to the endogenous TRIM28 protein containing a N-terminal KRAB-binding domain and C-terminal HP1, PHD, and BROMO domains; TRIM28^NFD^ with these C-terminal domains excised; and TRIM28^VPR^ with these C-terminal domains replaced by the synthetic transcriptional activator VPR (Fig. 1a). We applied a modified luciferase transfection assay into Neuro2a cells that our group has previously implemented^22^. In cultured Neuro2a cells, we simultaneously co-transfected: 1) a *luciferase* reporter plasmid under the control of a thymidine kinase (TK) promoter and possessing multiple ZFP189-recruiting DNA response elements (REs); 2) a plasmid overexpressing ZFP189, a representative KZFP that has previously been demonstrated to regulate this *luciferase* reporter gene^22^; and 3) a plasmid overexpressing one of our synthetic TRIM28 variants or a GFP plasmid control (Fig. 1b). Luciferase signal was calculated as relative light units (RLUs) as a proxy for gene transcription (Fig. 1c). Here, we saw no difference between luciferase signal in cells transfected with TRIM28^WT^ versus TRIM28^NFD^, but TRIM28^VPR^ did increase luciferase signal (p < 0.0001, one-way ANOVA with Bonferroni correction). This indicates that our TRIM28^VPR^ variant can behave as a gene activator at KZFP-regulated genes.

**Figure 1:**
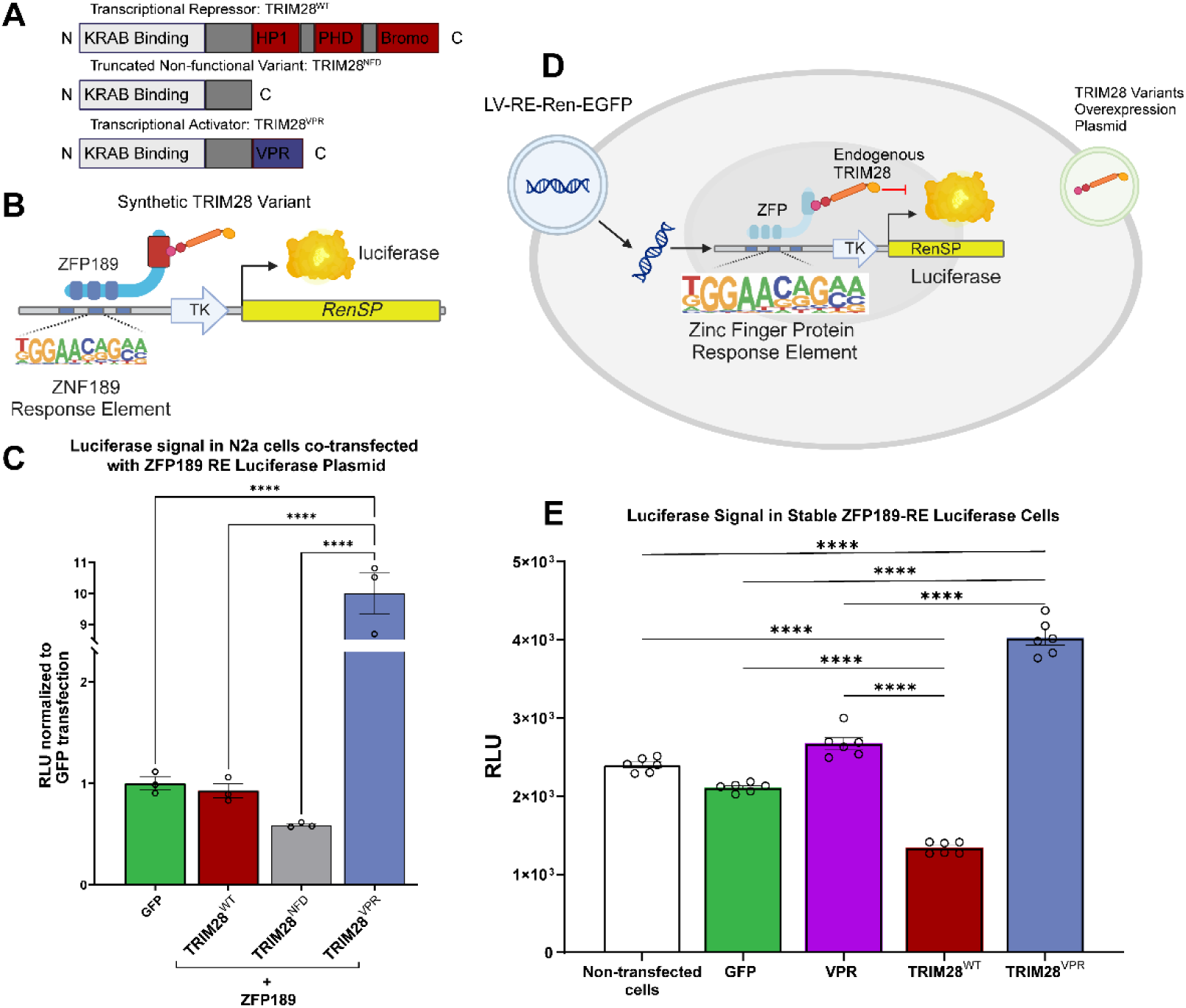
Synthetic TRIM28 variants bi-directionally control the expression of a KZFP-regulated gene. a) Schematic of novel synthetic TRIM28 variant proteins. b) Schematic of the luciferase reporter, which contains the KRAB-zinc finger protein (KZFP) ZFP189 response elements upstream of a thymidine kinase (TK) promoter and RenSP LightSwitch^TM^ luciferase gene. c) Co-transfection of Neuro2a (N2a) cells with separate overexpression vectors containing the luciferase reporter, ZFP189 (a representative KZFP), and our synthetic TRIM28 variants respectively shows TRIM28^VPR^ significantly upregulates luciferase signal, (p < 0.0001, one-way ANOVA with Bonferroni correction, n = 3 per group). d) Schematic of a stable N2a cell line that stably expresses the luciferase reporter (LV-RE-Ren-EGFP-N2a). These stable cells were then transfected once with a single overexpression vector containing synthetic TRIM28 variants. e) TRIM28^VPR^ significantly increases luciferase signal, while TRIM28^WT^ significantly decreases luciferase signal, compared to non-transfected control cells, or cells transfected with overexpression vectors for GFP or an isolated VPR domain (TRIM28^VPR^: p < 0.0001 versus non-transfected cells, GFP, and VPR. TRIM28^WT^: p < 0.0001 versus non-transfected cells, GFP, and VPR. One-way ANOVA with Bonferroni correction, n = 6 per group).

We also developed a Neuro2a cell line stably expressing the ZFP189-sensitive *luciferase* reporter (Fig. 1d). By transfecting these stable cells with TRIM28 variant expression plasmids, we observed that TRIM28^WT^ decreased luciferase signal compared to non-transfected cells, and against GFP control or isolated VPR domain transfections (p < 0.0001 for all). Conversely, we observed that TRIM28^VPR^ increased luciferase signal compared to non-transfected cells, and against GFP control or isolated VPR domain transfections (p < 0.0001 for all), indicating that TRIM28^WT^ and TRIM28^VPR^ produce opposite effects on KZFP-regulated gene transcription (Fig. 1e). Collectively, these *in vitro* studies indicate our synthetic TRIM28 variants can, as designed, specifically up- or down-regulate the expression of KZFP-regulated genes.

### Inverting TRIM28-mediated transcriptional control in prefrontal cortex causes social behavioral deficits in male and female mice

We next sought to characterize the *in vivo* behavioral consequences of dysregulating TRIM28-mediated transcriptional control in the PFC of male and female mice. We virally delivered our synthetic TRIM28 variants and GFP control via stereotaxic surgery to the PFC^31,32^ and, in distinct experimental cohorts, subjected test mice to social behavioral assays (Fig. 2a). As expected in a three-chamber social interaction task, when presented with choice between a sex and age-matched conspecific or an empty cage, all mice displayed a normal preference for social interaction (p < 0.0001 for all) (Fig. 2b). Notably, when presented with a novel versus familiar conspecific, all except HSV-TRIM28^VPR^ mice showed normal preference to interact with the novel conspecific (p < 0.0001, 0.0003, 0.0098 respectively) (Fig. 2c). Yet, mice receiving PFC HSV-TRIM28^VPR^ specifically did not show this preference for interacting with a novel sex and age-matched conspecific over a familiar conspecific (p > 0.05) (Fig. 2c). This lack of preference for novel social interaction cannot be explained by alterations in anxiety or locomotor phenotypes, as no viral treatment group displayed any deficits in elevated plus maze exploration, novelty suppressed feeding, or sucrose preference test (Supp. Fig. 1).

**Figure 2:**
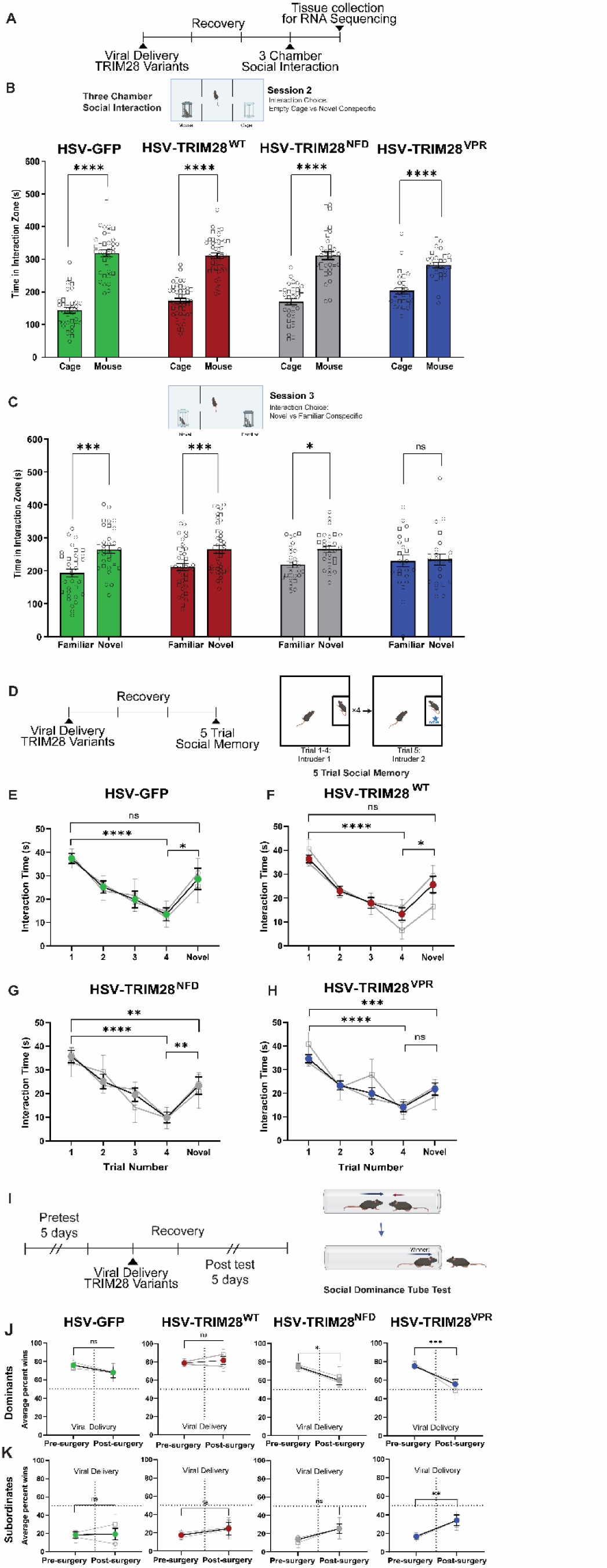
Disrupting PFC TRIM28 function disrupts social behavior. a) Timeline of three chamber social interaction and prefrontal cortex (PFC) tissue collection. Each vertical hash is a day (HSV-GFP: n = 10 females, 23 males; HSV-TRIM28^WT^: n = 10 females, 29 males; HSV-TRIM28^NFD^: n = 10 females, 18 males; HSV-TRIM28^VPR^: n = 6 females, 17 males). b) In Session II of three chamber social interaction (cage v. mouse), all viral conditions spend more time interacting with a novel conspecific than an empty cage (p < 0.0001, two-way ANOVA with Bonferroni correction). c) In Session III of three chamber social interaction (familiar v. novel), HSV-TRIM28^VPR^ ablates the preference for socializing with a novel conspecific over a familiar one (p > 0.05, two-way ANOVA with Bonferroni correction), while this preference is preserved in HSV-GFP, HSV-TRIM28^WT^, and HSV-TRIM28^NFD^-treated mice (p < 0.0001, 0.0003, 0.0098 respectively, two-way mixed effects ANOVA with Bonferroni correction). d) Timeline and experimental design for five trial social memory (HSV-GFP: n = 8 females, 11 males; HSV-TRIM28^WT^: n = 8 females, 19 males; HSV-TRIM28^NFD^: n = 8 females, 19 males; HSV-TRIM28^VPR^: n = 8 females, 27 males). e-g) Mice receiving HSV-GFP, HSV-TRIM28^WT^, or HSV-TRIM28^NFD^ habituate to a repeated social partner and renew interest for a novel partner (Trial 1 v. 4: p < 0.0001 for all, Trial 4 v. novel: p = 0.0123, 0.0309, 0.0024 respectively, two-way ANOVA with Bonferroni correction). Only HSV-TRIM28^NFD^-treated mice spend significantly less time interacting with the novel interaction target than with the initial interaction target (Trial 1 v. novel, p = 0.0098, two-way ANOVA with Bonferroni correction). h) Mice receiving HSV-TRIM28^VPR^ habituate to a repeated social partner, but do not renew interest for a novel partner (Trial 1 v. 4: p < 0.0001, Trial 4 v. novel, p > 0.05, two-way ANOVA with Bonferroni correction). These mice also spend significantly less time interacting with the novel interaction target than with the initial interaction target (Trial 1 v. novel, p = 0.0003, two-way ANOVA with Bonferroni correction). i) Timeline and experimental design for social dominance tube test. j) Comparing average pretest win rate to average posttest win rate, dominant mice who received HSV-TRIM28^NFD^ (n = 6 females, 11 males) and -TRIM28^VPR^ (n = 6 females, 12 males) had significantly decreased win rates in the posttest (p = 0.0092 and p = 0.0002 respectively, two-way ANOVA with Bonferroni correction). Receiving HSV-GFP (n = 6 females, 6 males) or -TRIM28^WT^ (n = 6 females, 6 males) did not significantly alter dominant mouse win rates (p> 0.05, two-way ANOVA with Bonferroni correction). k) Comparing average pretest win rate to average posttest win rate, subordinate mice who received HSV-GFP (n = 6 females, 6 males), -TRIM28^WT^ (n = 6 females, 6 males), or -TRIM28^NFD^ (n = 6 females, 11 males) did not have significantly different win rates (p > 0.05, two-way ANOVA with Bonferroni correction), while subordinate mice who received HSV-TRIM28^VPR^ (n = 6 females, 12 males) had a significant increase in win rate (p = 0.0099, two-way ANOVA with Bonferroni correction). Squares indicate females; circles indicate males.

We also aimed to confirm whether these social deficits could be explained by deficits in novelty recognition in general, or whether this was a specific social recognition deficit. We performed a five-trial social memory test (Fig. 2d), where experimental mice are subjected to four sequential sessions with the same age and sex-match conspecific, followed by a fifth session with a novel conspecific^22,33^. Typically, mice will spend decreasing amounts of time interacting with the same conspecific through sessions 1-4, and renew social interaction behaviors during session 5. Mice receiving HSV-TRIM28^WT^, -TRIM28^NFD^, and -GFP all displayed this characteristic pattern of behavior (Fig. 2e-h). However, mice receiving HSV-TRIM28^VPR^ did not show a significantly renewed interest in interacting with the novel mouse in session 5, suggesting the salience of novel social interaction with the new conspecific was diminished (p > 0.05, 2-way ANOVA with Bonferroni correction) (Fig. 2h). Yet, the progressive decrease in interaction time through sessions 1-4 for the HSV-TRIM28^VPR^ mice does suggest that the capacity for social memory is intact (p < 0.0001).

To more completely characterize the nature of TRIM28^VPR^-mediated social behavioral deficits, we performed repeated social dominance tube tests (Fig. 2i). Our group has previously employed this task to characterize how viral delivery of synthetic TFs affect awareness and participation in social hierachies^22^. Conventionally, dominance is defined as consistently winning the majority of tube tests against cage mates, while subordinance reflects consistent losses against cage mates^34,35^. Here, we allowed cages of five sex-matched mice to establish a social hierarchy, then delivered HSV-GFP, -TRIM28^WT^, -TRIM28^NFD^, or -TRIM28^VPR^ to the PFC of both dominant and subordinate mice, and finally retested the cage’s social hierarchy for five days post-surgery. Intermediate ranked mice received intra-PFC HSV-GFP to match surgical experience to the test mice. We elected to analyze results pre-vs post-surgery between viral conditions separately between dominant mice and subordinate mice.

As expected, dominant and subordinate mice receiving HSV-GFP and HSV-TRIM28^WT^ remained dominant and subordinate respectively, with win rates remaining significantly different from random chance post-surgery (Dominants: p = 0.0171, 0.0010 respectively, Subordinates: p = 0.0112. 0.0107 respectively, Wilcoxon signed-rank test) (Fig. 2j, k). Mice receiving HSV-TRIM28^NFD^ did not completely maintain their social hierarchy positions post-surgery, as dominant mice became significantly less dominant (p = 0.0092, two-way ANOVA with Bonferroni correction), though they still won more often than random chance (p = 0.0215, Wilcoxon signed-rank test). Yet, subordinate mice receiving HSV-TRIM28^NFD^ remained similarly subordinate (p > 0.05, two-way ANOVA with Bonferroni correction) and maintained a lesser win rate compared against random chance, indicating the maintained awareness of their hierarchical position (p = 0.0366, Wilcoxon signed-rank test) (Fig. 2j, k). However, HSV-TRIM28^VPR^ caused both dominant and subordinate mice to revert towards a random chance of winning in the binary outcome of the tube-test (p > 0.05, Wilcoxon signed-rank test) (Fig. 2j, k). Further, dominant and subordinate mice receiving HSV-TRIM28^VPR^ all performed significantly different post-surgery compared to their pre-surgery hierarchical position (p = 0.0002, 0.0099 respectively, two-way ANOVA with Bonferroni correction). This indicates inversion of endogenous PFC TRIM28 molecular function is sufficient to remove the mouse’s capacity to participate in social hierarchies and/or remove their capacity for performing cooperative social strategies required to consistently resolve a social conflict, here modeled in a head-to-head tube test.

Finally, we tested novel object recognition to determine if this deficit in novel social interaction could be explained by deficits in recognition of non-social stimuli. We applied a previously validated Y-maze novel object recognition paradigm^36,37^ and discovered mice in all treatment groups displayed identical performance on the Y-maze spontaneous alternation task as well as retained ability to recognize novel objects (Supp. Fig. 2). These data suggest that HSV-TRIM28^VPR^ uniquely disrupts aspects of social behaviors, but not other behaviors relying on general non-social recognition.

### TRIM28^VPR^ derepresses transposable elements in the prefrontal cortex

In order to understand the TRIM28-driven brain molecular mechanisms associated with the observed social deficits, we performed bulk RNAseq at day four post-transduction on individual microdissected PFC tissue from individual male and female brains transduced with each of our synthetic TRIM28 variants. This timeline was selected to reflect the timing of emergence of the behavioral changes observed above (Fig. 2). Given the growing body of literature linking KZFPs to silencing mechanisms of TEs^22,23,26,28,38^, we hypothesized that inversion of TRIM28 gene-regulatory function achieved by overexpression of HSV-TRIM28^VPR^ would lead to widespread dysregulation of such TEs. Thus, we utilized a modified RNAseq analytical pipeline that included analyses for TEs, which are typically excluded in RNAseq analyses due to multiple read alignment^39,40^. All differentially expressed genes (DEGs) and differentially expressed TEs (DETEs) were calculated relative to the HSV-GFP viral control condition. We observed that

HSV-TRIM28^WT^ produced up- and down-regulation of DEGs, and, as expected, did not produce many DETEs compared to the GFP control (Fig. 3a; full DEG and DETE gene list in Supp. Table 1). In contrast, HSV-TRIM28^NFD^ produced more upregulated DETEs (Fig. 3b; full DEG and DETE gene list in Supp. Table 2), pointing to the possibility for a potential dominant negative function of TRIM28^NFD^ competing with endogenous PFC TRIM28 and derepressing TEs. Most strikingly, HSV-TRIM28^VPR^ asymmetrically drove robust upregulation of DETEs (Fig. 3c; full DEG and DETE gene list in Supp. Table 3), indicating that the additional gene-activating function of the VPR moiety on TRIM28^VPR^ serves to actively drive TE escape.

**Figure 3:**
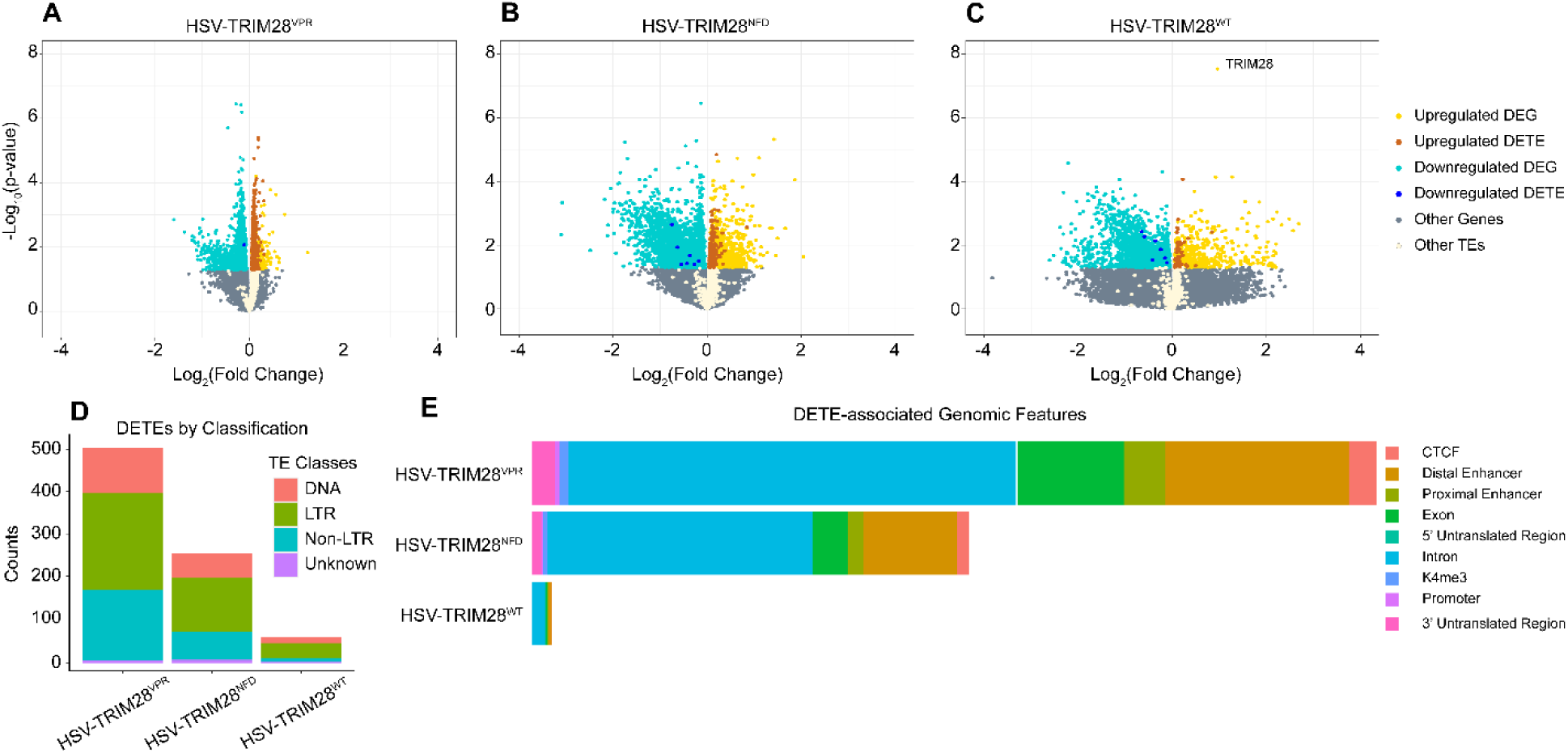
Disrupting PFC TRIM28 function drives the release of transposable elements. a-c) Volcano plots of all gene and transposable element transcripts induced by HSV-TRIM28^VPR^, HSV-TRIM28^NFD^, and HSV-TRIM28^WT^ normalized to the GFP control condition (HSV-GFP: 9 females, 9 males; HSV-TRIM28^WT^: 10 females, 10 males; HSV-TRIM28^NFD^: 8 females, 9 males; HSV-TRIM28^VPR^: 10 females, 9 males). Genes (DEGs) and TEs (DETEs) were considered statistically significant if p < 0.05. HSV-TRIM28^VPR^ induced many more DETEs compared to other TRIM28 variants. d) HSV-TRIM28^VPR^ and HSV-TRIM28^NFD^ induced activation of all classes of TEs, and TEs of the non-LTR class were disproportionately activated (Chi-square goodness of fit test, p < 0.0001 for both). e) HSV-TRIM28^VPR^ and HSV-TRIM28^NFD^ activated TEs that were predominantly derived from intronic and distal enhancer genomic features.

We next assessed the subcategories of DETEs dysregulated by each of our synthetic TRIM28 variants. Irrespective of direction of regulation, HSV-TRIM28^WT^ produced 61 DETEs compared to the GFP control, while HSV-TRIM28^NFD^ produced 259 DETEs and HSV-TRIM28^VPR^ produced 510 DETEs (Fig. 3d). Further, the distribution of LTR, non-LTR, and DNA transposons differed across viral treatment conditions (Supp. Table 4). In all TRIM28 variant conditions, TEs in the LTR class account for approximately half, and DNA transposons account for approximately 20% of all DETEs detected. However, non-LTR class TEs expanded from 13% of DETEs in HSV-TRIM28^WT^ to 26% of DETEs in HSV-TRIM28^NFD^ and 33% of DETEs in HSV-TRIM28^VPR^, indicating the non-LTR class of TEs were disproportionately derepressed by our synthetic TRIM28 variants compared to HSV-TRIM28^WT^ (Chi-square goodness of fit, p < 0.0001 for HSV-TRIM28^NFD^ and HSV-TRIM28^VPR^). Additionally, we annotated the genomic features with which these DETEs overlap. All three TRIM28 variants affected DETEs originating from intronic and distal enhancer genomic regions, accounting for approximately 80% of all annotated genomic features (Fig. 3e, Supp. Table 5). These transcriptomic data suggest that inverted TRIM28 function in PFC nerve cells derepresses primarily non-LTR TEs residing within genomic distal enhancer and intronic regions.

### Derepression of transposable elements is associated with downregulation of immune genes

We next sought to explore the differences in canonical gene transcription produced by our synthetic TRIM28 variants. We first applied rank-rank hypergeometric overlap (RRHO) analyses to assess the broad congruence of gene-expression patterns between our synthetic TRIM28 variants. RRHO captures threshold-free similarities and differences between gene lists^41^. Using RRHO, we observed that the collective gene expression profile affected by HSV-TRIM28^VPR^ was more similar to HSV-TRIM28^NFD^ than to HSV-TRIM28^WT^ (Fig. 4a, b). Further, HSV-TRIM28^NFD^ produced an intermediate transcriptional profile that was more broadly similar to HSV-TRIM28^WT^ than to HSV-TRIM28^VPR^ (Fig. 4b, c), pointing to a gradation in the concordance of PFC transcriptional profile that is stratified across TRIM28 normal function (HSV-TRIM28^WT^), non-function (HSV-TRIM28^NFD^), and anti-function (HSV-TRIM28^VPR^). This also mirrors the gradation of behavioral effects observed in Figure 2.

**Figure 4:**
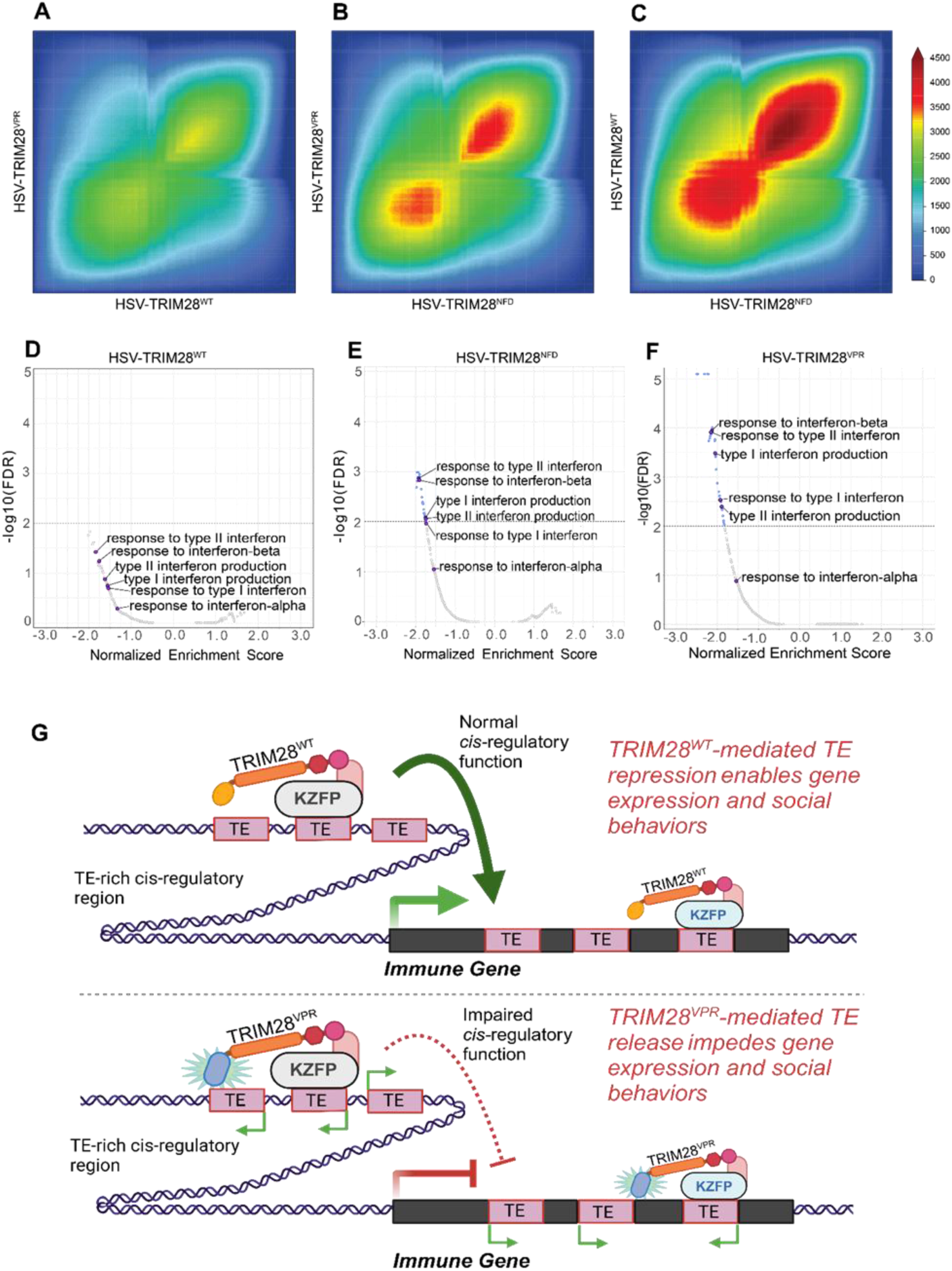
Disrupting PFC TRIM28 function impairs immune gene expression. a-c) Rank-rank hypergeometric overlap graphs demonstrate threshold free comparisons between gene lists. Transcription induced by HSV-TRIM28^VPR^ is more similar to HSV-TRIM28^NFD^ than HSV-TRIM28^WT^. d-f) Volcano plots showing gene set enrichment analysis terms generated from all canonical genes in HSV-TRIM28^WT^, -TRIM28^NFD^, and -TRIM28^VPR^, all normalized to HSV-GFP-treated mice. Interferon-related gene ontology terms are specifically labeled. A significance cutoff of FDR < 0.01 was applied. Interferon-related ontology terms were significantly downregulated by HSV-TRIM28^VPR^ to a greater extent than HSV-TRIM28^NFD^, and were not affected by HSV-TRIM28^WT^. g) Cartoon depicting the potential molecular mechanism linking PFC TRIM28 function and dysfunction to transposable elements, gene expression, and social behaviors.

To characterize the nature of affected genes in each condition, we performed gene set enrichment analysis (GSEA) of total DEGs produced by each TRIM28 variant (Fig. 4d-f). To improve specificity of ontology terms while preserving an exploratory approach, the maximum number of gene IDs per category was limited to 300, and only terms with a false discovery rate (FDR) < 0.01 were designated significant. In the GSEA, which accounts for both the directionality and magnitude of expression change, we observed that while all three TRIM28 variants primarily display downregulated gene ontology terms, the magnitude of this downregulation mirrors the gradation of effect observed in both TE upregulation (Fig. 3) and behavioral dysregulation (Fig. 2). In particular, we observed that interferon-related terms adhered to the same pattern of downregulation, with HSV-TRIM28^WT^ inducing a non-significant downregulation (Fig. 4d), HSV-TRIM28^NFD^ a moderate but significant downregulation (Fig. 4e), and HSV-TRIM28^VPR^ a greater significant downregulation of interferon-related terms (Fig. 4f). Upon consulting our DEG tables, this is directly recapitulated in the expression levels of interferon genes across our TRIM28 variants (Supp. Fig. 3). Only HSV-TRIM28^VPR^ caused significant downregulation of interferon beta and gamma, while HSV-TRIM28^NFD^ caused downregulation of interferon alpha-B and HSV-TRIM28^WT^ did not downregulate any interferon cytokines (Supp. Fig. 3). Accounting for the growing body of literature linking immune functions to social behaviors^21,42,43^, including from our group^22^, this represents a potential molecular mechanism explaining the HSV-TRIM28^VPR^-driven behavioral deficits. We illustrate the potential mechanistic relationship between KZFP-TRIM28 gene regulatory control, genomic TEs, immune gene function, and social behaviors in the cartoon represented in Figure 4g.

As described in Figure 3, our synthetic TRIM28 variants induced activation of TEs, and these DETEs primarily originated from genomic TE sequences localized within intronic and distal enhancer genomic features. Previous studies investigating cis-regulatory element chromatin looping suggest TE-derived cis-regulatory elements typically exert transcriptional control on genomically distant target genes^44,45^. Other studies note such distant TE-derived enhancers are located in intergenic or intronic TE-rich regions^46^, suggesting that HSV-TRIM28^VPR^-mediated escape of such enhancer TEs may disrupt cis-regulation of target genes and causally contribute to the transcriptional divergence across TRIM28 variants. Thus, we also interrogated genes with noted physical association with DETEs as predicted in EnhancerAtlas^47^. We identified TE-associated DEGs by applying a p-value cutoff of 0.05 to the DEGs with a predicted association to any DETE in our RNAseq datasets. Using this method of filtering the gene expression tables, we observed 142 TE-associated DEGs in the HSV-TRIM28^WT^ condition, 2495 in the HSV-TRIM28^NFD^ condition, and 1665 in the HSV-TRIM28^VPR^ condition. These TE-associated DEGs are majority downregulated, with 124 downregulated in HSV-TRIM28^WT^, 1998 downregulated in HSV-TRIM28^NFD^, and 1295 downregulated in HSV-TRIM28^VPR^. Applying GSEA analysis to these TE-associated DEGs (Supp. Fig. 4), we observe a broadly similar pattern to the unfiltered DEG analysis (Fig. 4d-f). Thus, we posit that the overall transcriptional profile characterized by downregulation of immune-related genes may be predominantly driven by TE-associated genes (Supp. Fig. 4).

### Repletion of PFC immune function rescues TRIM28^VPR^-driven social deficits

To determine whether the observed social deficits were causally produced by HSV-TRIM28^VPR^-driven deficiencies in PFC immune function, we tested if treating HSV-TRIM28^VPR^ mice with exogenous interferon could restore normal social behavior. We co-delivered HSV-TRIM28^VPR^ with a mixture of 10 ng each of interferon beta and gamma (IFN) or saline vehicle (VEH) and completed three-chamber social interaction on day four post-surgery, mirroring the timeline used in Figure 2 (Fig. 6a). We observed that transduction with HSV-TRIM28^VPR^ + VEH recapitulated the social behavior deficit in session three as previously observed, with mice displaying no preference for novel social interaction (Fig. 6b, p > 0.05, two-way ANOVA with Bonferroni correction). However, HSV-TRIM28^VPR^ + IFN mice restored a preference for novel social interaction in this session (Fig. 6c, p = 0.0071, two-way ANOVA with Bonferroni correction).

Interestingly, in session two, HSV-TRIM28^VPR^ + IFN mice displayed further increased sociability compared to HSV-TRIM28^VPR^ + VEH, spending increased total time spent socializing with the target mouse (Fig. 6b, p = 0.0409, two-way ANOVA with Bonferroni correction). Lastly, we confirmed that administration of exogenous IFN did not impede HSV-mediated transduction of PFC neurons (Supp. Fig. 5). These data indicate that disruption of endogenous immune functions is directly responsible for the observed HSV-TRIM28^VPR^-mediated social deficits and that these behavioral deficits are reversible.

## Discussion

Here we present research to demonstrate that TRIM28 functions in mouse PFC to stabilize genomic TEs, enable appropriate immune gene expression, and facilitate complex social behaviors characterized by awareness and interest in social novelty. By inverting normal repressive TRIM28 function using synthetic HSV-TRIM28^VPR^, we demonstrated that mice developed social deficits characterized by loss of preference for novel social interaction in three chamber social interaction and five-trial social memory tasks and a diminished capacity to engage in complex social hierarchies in the social dominance tube test (Fig. 2), without producing any non-social behavioral deficits. Further, we observed HSV-TRIM28^VPR^ caused substantial activation of TEs and downregulation of genes involved in immune ontology terms including response to type 1 and type 2 interferon (Fig. 3, Fig. 4). HSV-TRIM28^VPR^-driven social deficits were reversible by exogenous supplementation of these interferon cytokines, indicating that these social deficits were causally produced by impairments in PFC immune signaling.

These data are consistent with a growing body of literature linking social behaviors with brain adaptive immune functions^6–8,23,25,48–55^. Here, we discover that KZFP/TRIM28-mediated genomic stability of cortical TEs is a key molecular mediator that links brain immune gene expression and complex social behaviors, suggesting that TE silencing in this brain region is not solely established at singular developmental timepoint, but rather is constantly maintained by the persistent functions of TRIM28 and related molecular processes. Thus, the conditions in which KZFPs, TRIM28, and TEs become dysregulated in brain may contribute to our understanding of social dysfunction that characterizes many neuropsychiatric syndromes.

Interestingly, we observed HSV-TRIM28^VPR^-mediated downregulation of immune genes is highly enriched for DEGs associated with activated TEs (Supp. Fig. 4). Given the existing literature linking TE-rich genomic regions with cis-regulatory mechanisms^14–19^, we posit that these TRIM28-controlled genomic loci may regulate immune genes, particularly interferons, through cis-regulatory mechanisms that require stabilization and repression of TEs to maintain appropriate expression. Notably, our TE-associated DEG analysis revealed that not only do TEs residing in intronic and distal enhancer regions of the genome account for over 70% of all TE origins detected in our data, but they also account for over 90% of all TE-associated DEGs. This recapitulates existing literature suggesting that cis-regulatory mechanisms evolved by co-opting specifically intronic and intergenic TE-rich regions and further supports our hypothesized mechanism (Fig. 4g)^46^.

While the immune-related DEG profile of HSV-TRIM28^VPR^- and HSV-TRIM28^NFD^-treated mice is consistent with a decrease in inflammatory signaling (Fig. 4e-f), the social behavioral profile is similar to that of a sickness-like or inflammation-induced phenotype, which is characterized by a decreased social novelty preference and a disruption to social hierarchy^56–58^. However, mice lacking adaptive immunity (SCID mice) or the IFNγ gene also display a lack of social approach reversible by introduction of lymphocytes from IFNγ-competent mice^21^, similar to how interferon repletion prevented the deficits induced by HSV-TRIM28^VPR^ (Fig. 5). This suggests that a balance of immune system activity is required to maintain a typical social behavioral profile.

**Figure 5:**
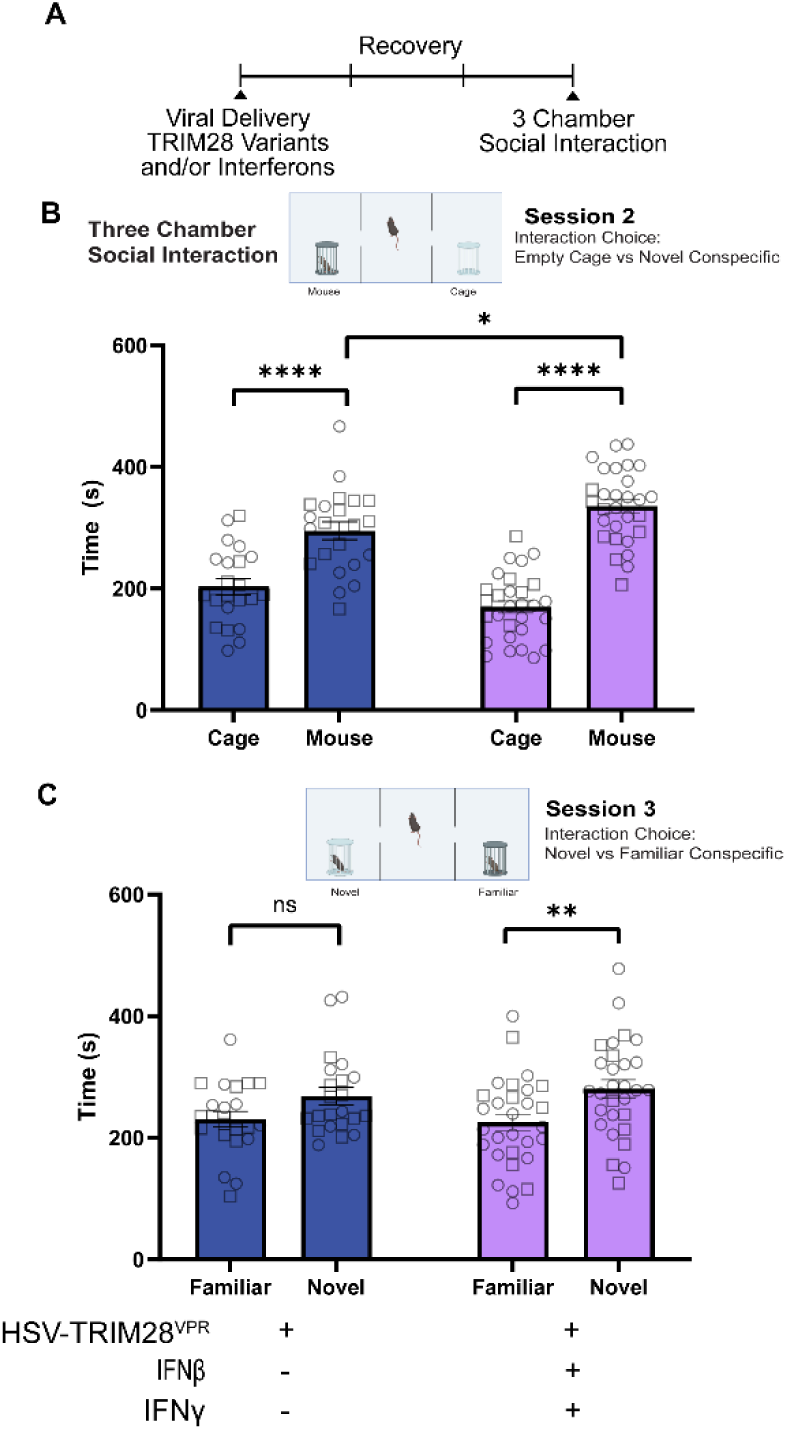
Repletion of PFC immune function rescues TRIM28^VPR^-driven social deficits. a) Timeline of viral delivery and 3 Chamber Social Interaction testing. Each hash represents one day (HSV-TRIM28^VPR^ + VEH: n = 10 females, 12 males; HSV-TRIM28^VPR^ + IFN: n = 9 females, 20 males). b-c) Co-administration of HSV-TRIM28^VPR^ with exogenous IFNβ and IFNγ increases time spent socializing with a novel age and sex-matched conspecific in session 2 of 3 chamber social interaction (p = 0.0409, two-way ANOVA with Bonferroni correction). Both HSV-TRIM28^VPR^ + IFN and HSV-TRIM28^VPR^ + VEH-treated groups displayed preference for social interaction over the empty cage (p < 0.0001, two-way ANOVA with Bonferroni correction). In session 3, HSV-TRIM28^VPR^ + VEH displayed a lack of preference for novel social interaction, while HSV-TRIM28^VPR^ + IFN restored preference for novel social interaction (p > 0.05, = 0.0077 respectively, two-way ANOVA with Bonferroni correction). Squares indicate females; circles indicate males.

All experiments presented in this study were completed with both male and female mice. In our behavioral studies, we initially included sex as a variable in multi-way ANOVA statistical analysis to account for the possibility of detecting a sex difference in the behavioral consequences of dysregulation of TRIM28 transcription. However, we elected to combine all male and female data as there was no significant difference between sexes in any assay measured (Three-way ANOVA, main effect for sex p > 0.05). The absence of a detectable sex difference in our social behavioral data supports the notion that TRIM28-mediated PFC molecular mechanisms similarly govern complex social behavior in both male and female mice, irrespective of well-noted baseline sex differences in the rodent social behaviors^59,60^.

Notably, chromatin modifications and TE escape similar to synthetic induction via HSV-TRIM28^VPR^ have been linked to ASD and ASD-like phenotypes in human populations and in animal models, particularly those with social behavior deficits^61–70^. Heterozygous deletions or conditional knockouts of *TSHZ3*, a zinc finger protein, in mice cause changes in social and repetitive behaviors similar to individuals with ASD^62,68^. In particular, *TSHZ3* conditional knockout in cortical projection neurons, such as those targeted by our PFC-delivered HSV, specifically impacts social, but not repetitive, behaviors^68^. Additionally, the inhibition of brain chromatin modifying enzymes such as HDACs has been shown to both induce and rescue social behaviors in different mouse models of ASD^61,63–65^. Further, prenatal exposure to the non-specific HDAC inhibitor valproic acid causes the release of endogenous retroviruses, which persists across multiple generations of mice and is associated with social deficits that also persist across generations^64^. TE transposition has been linked to social behaviors in canine studies, with the direction of effects dependent upon the genes affected by TE insertion^70^. In several human sequencing studies, insertions of TEs have been linked to ASD, and these are particularly prevalent in intronic and intergenic regions, often enriched in known ASD risk genes^66,67,69^. Thus, our HSV-TRIM28^VPR^-induced TE derepression adds to the existing literature on social behavior disruptions linked to TE transcription.

Previously, our group has shown that a particular member of the KZFP family, ZFP189, modulates social behavior while tuning TE-dependent, immune-related neurobiological mechanisms^22^. Here, we utilize synthetic TRIM28 variants to induce transcriptional dysregulation via the collective function of the KZFP family of proteins. We observed consistency in results, as inverting the function of either ZFP189 or TRIM28 induced social behavioral deficits while sparing non-social behaviors. This points to PFC KZFPs beyond just ZFP189 as involved in regulating social behaviors.

In sum, we present a novel synthetic biology approach to collectively dysregulate the transcriptional control of the KZFP family of proteins within the PFC via re-engineering of their key binding partner TRIM28. In this manner, we observe that intra-PFC function of TRIM28 is necessary to maintain normal social behaviors, particularly higher-order social cognition involved in the salience of novel social interactions. Further, we showed that the gene transcriptional changes caused by inversion of TRIM28 transcriptional control are characterized by derepression of TEs and disruption of immune-related genes modulated by associated TE-rich genomic regions. This work further clarifies the brain molecular link between PFC KZFP function, TE stability, and immune expression, all of which form a molecular regulatory axis to enable complex social behaviors necessary for group living.

## Methods

### Subjects

Male and female C57BL/6 J mice (8 to 10 weeks old) from Jackson Laboratories were used. Mice were group housed (5 mice/cage) on a 12-hour light/dark cycle (lights on at 6am/off at 6 pm) with ad libitum access to food and water. All mice were used in accordance with protocols approved by the Institutional Care and Use Committees at Virginia Commonwealth University School of Medicine.

### Viral packaging

We de novo synthesized TRIM28^NFD^, TRIM28^WT^, and TRIM28^VPR^ and sub-cloned each variant into HSV expression plasmids via ThermoFisher Scientific gateway LR Clonase II cloning reaction and Gateway LR Clonase II Enzyme mix kit (catalog number 11791-020 and 11971-100). Colonies were Maxi-prepped (Qiagen Cat # 12163) and shipped to the Gene Delivery Technology Core at Massachusetts General Hospital for HSV packaging. Once packaged, aliquots were made and stored in −80 °C to be used in viral gene transfer through stereotaxic surgery.

### Viral gene transfer

Stereotaxic surgeries targeting the PFC were performed as previously described^31,32^. Mice were anesthetized with I.P. injection of ketamine (100 mg/kg) and xylazine (10 mg/kg) dissolved in sterile saline solution. Mice were then placed in a small-animal stereotaxic device (Kopf Instruments) and the skull surface was exposed. 33-gauge needles (Hamilton) were utilized to infuse 1.0 μL of virus at a rate of 0.2 μL/min followed by a 5-min rest period to prevent backflow. For the interferon repletion experiment, HSV-TRIM28^VPR^ was co-delivered with a mixture of 10 ng each of interferon beta and gamma (ThermoFischer) (IFN) or saline vehicle (VEH). The following coordinates were used to target the PFC: Bregma: anterior-posterior: +1.85 mm, medial-lateral ±0.75 mm, dorsal-ventral −2.5 mm, 15° angle.

### N2a Cell culture

*Mus musculus* Neuro-2a (N2a; ATCC^®^ CCL-131™) neuroblast cell culture were maintained in 1:1 EMEM enriched / EMEM growth medium mixture (Quality Biological, #112-039-101; ATCC, #30 2003 or Corning, #10-009-CV) with 5% FBS (HyClone, #SH30071.03IH30-45) and 100 U/ml Penicillin Streptomycin (Gibco, #15140122) in a 37 °C and 5% CO_2_ Thermo Scientific HERAcell-150i CO_2_ incubator. N2a cells were split twice weekly at 1:6 up to passage ∼50 as previously described^22^.

### ZNF189RE-RenSP pLV[Exp]-EGFP:T2A:Puro Cloning and lentivirus packing

The ZNF189 response element (RE)-basal TK promoter-RenSP luciferase gene from our previous ZNF189RE-RenSP luciferase reporter vector^22^ was synthesized and subcloned into a Mammalian Gene Expression Lentiviral Vector (pLV[Exp]-EGFP:T2A:Puro) and packaged in lentivirus at a titer of 4.82 ×10^8^ TU/ml by VectorBuilder. The vector ID is VB230720-1406wne, which can be used to retrieve detailed information about the vector on vectorbuilder.com.

### ZNF189RE-RenSP-N2a stable cell pool

Low passage N2a cells (p5.5) were plated in a 6-well plate (3×10^5^ cells per well) and were grown until 70% confluent at the time of transduction. Following the Lentivirus In Vitro Applications User Instructions (Version 2.0, 2022-01-07) from VectorBuilder, cells were infected with the virus at multiplicity of infection (MOI) = 2, 3, and 5 in 1ml of medium. When the vector EGFP expression was visible under a fluorescence microscopy (Bio-Rad ZOE^TM^ Fluorescent Cell Imager) from day 2 of post-infection, puromycin selection (1.5ug/ml in 3ml of medium per well) was applied for 4 days until uninfected cells were killed. Transduced cells with MOI of 2, 3, and 5 were mixed together as a ZNF189RE-RenSP-N2a stable pool and maintained in medium containing 2ug/ml puromycin.

### Transfection and luciferase reporter assay

The day before transfection, approximately 1.5-2×10^4^ cells in 100ul medium per well (N2a or RE-RenSP-N2a stable pool) were seeded into 96-well clear bottom microplates with white walls (Corning, # 3610). Wells reached roughly 80-90% confluent in 24 hours. Using normal N2a cells, we applied the Effectene Transfection Reagent (Qiagen # 301427) to co-transfect the reporter plasmid DNA of ZNF189 RE (25ng) with our synthetic TRIM28 variants and an unmodified ZFP189 expression plasmid DNA (100-120ng with equal molar weight). In our RE-RenSP-N2a stable pool, we performed transfection only with TRIM28 variant expression vectors using the same transfection protocol. An unmodified GFP empty expression vector (p1005gw Δ*CCDB*) and an expression plasmid containing an untethered VPR domain were used as background controls.

On day 3 post-transfection, using a BMG Labtech POLARstar Omega Microplate Reader, the GFP Fluorescent (FI) was measured for equal transfection efficiency confirmation and normalization, followed by relative luminescence unit (RLU) using Renilla luciferase assay system (Promega, #E2820). Data was analyzed using GraphPad Prism 10 software.

### Behavioral testing

Behavioral analyses were performed automatically by video tracking software (Ethovision Noldus)^71^. All behavioral tests were performed in a specified behavioral suite under red light.

### Three-chamber social interaction test

The three-chamber test was used to assess sociability and interest in social novelty or social discrimination. The testing arena consisted of three adjacent chambers (each 41 × 21 × 41 cm) separated by two clear plastic dividers and connected by open doorways (5 × 9 cm). The test consisted of three 10-min sessions. The subject mouse begins each session in the middle chamber. In the first session, subject mice were habituated to the arena and allowed free investigation of the three chambers. In the subsequent sociability session, a novel C57BL/6J sex and age-matched conspecific (target mouse 1) was placed in a cylindrical cage (20 cm height × 10 cm diameter solid bottom; with clear bars spaced 2 cm apart) in one of the side chambers and an identical cage was placed in the opposite side chamber. In the social novelty session, the empty cage was replaced by target mouse 1 in a fresh cage. A second novel C57BL/6J sex and age-matched conspecific (target mouse 2) was placed at the previous position of target mouse 1 in a new cylindrical cage. The chamber and cylindrical cages were thoroughly cleaned with 70% EtOH between animals and before the first animal. Time spent in each chamber was recorded. Sociability was measured by comparing the time spent in the chamber with the novel mouse vs an empty cup. Social novelty was measured by comparing the time spent in the chamber with a novel vs familiar mouse.

### Five-trial social memory test

Five-trial social memory was tested as previously described^22,33^ to determine ability to recognize novel versus familiar animals. Subject mice were placed into the open arena (43 × 43 × 43 cm) with an empty wire cage (10 × 5 × 30 cm) at one side (interaction zone) and given 2.5 min of habituation to explore the arena before being returned to their home cage. A novel C57BL/6J sex and age-matched conspecific was then placed within the wire cage (interaction zone), and subject mice were placed back into the open arena for four subsequent 2.5 min trials with a 10 min inter-trial interval. In trial 5, a novel C57BL/6J sex and age-matched conspecific was placed within the interaction cage to measure dishabituation. Time spent in the interaction zone for the first minute of each trial was measured.

### Social dominance tube test

Animal social dominance was tested as previously described^22,34,35^ in a transparent Plexiglas tube measuring 30.5 cm in length and 3 cm diameter, a size sufficient to permit one subject mouse to pass through without being able to reverse direction. The tube was set on a plastic table in the designated behavioral suite and trials were manually recorded by a researcher blind to experimental groups. Animals were placed at opposite ends of the tube and released. A subject was declared the “winner” when its opponent backed out of the tube, with all 4 paws outside of the tube. The maximum test time allowed was 2 min. For 5 days, baseline social hierarchy was determined by once-daily tube tests for all animals within a five-mouse cage in a randomized order. In each cage, the most dominant and most subordinate mice received intra-PFC HSV-TRIM28 variants whereas the remaining cage-mates received intra-PFC HSV-GFP. For the following 5 days, the tube tests were repeated to determine post-surgery social hierarchy. Social dominance was measured by calculating the percentage of wins in the tube test (number of wins/number of tests * 100).

### Elevated plus maze

The EPM apparatus is constructed of black Plexiglas and consists of two open arms (33 × 6 cm) and two closed arms (33 × 9.5 × 20 cm) facing connected by a central platform (5 × 7 cm)^72^. The maze was elevated 63 cm above the floor. C57BL/6 J mice (45 mice across two cohorts following 3 chamber social interaction testing) were placed individually in the right-side closed arm facing the center of the plus-maze. Placement of all four paws into an arm was registered as an entry in the respective arm. The time spent in each arm was recorded during the 5 min EPM test. The platform of the maze was cleaned with 70% EtOH following each trial and before the first trial. The percentage of time spent in the open arms was calculated (time spent in open arms/300 s * 100 = % time spent in open arms).

### Novelty suppressed feeding

The novelty suppressed feeding arena consisted of a large rat cage (30 x 19 x 39 cm) containing woodchip bedding and a single pellet of normal chow. Following 8 hours of food restriction beginning at lights on, mice were individually placed in the arena for 5 minutes. Latency to feed was manually scored by a researcher blind to treatment. If a test mouse did not approach the food and begin to feed by the end of the 10-minute trial, the mouse was returned to its home cage and a latency of 10 minutes was recorded.

### Novel object recognition test

Novel object recognition was tested as previously described^36,37^. 24 hrs prior to the test day, subject mice were allowed to freely explore and habituate to the Y-maze arena for 10 minutes. Correct alternations were scored based on whether a mouse explored all three arms of the maze before repeating an arm. On the test day, subject mice underwent two test trials with a 10-minute inter-trial interval. The first trial presents subject mice with two identical object copies in two arms of the Y-maze arena. Following 5 minutes of exploration, subject mice were returned to their home cage. During the inter-trial interval, the arena and both interaction objects were thoroughly cleaned with 70% EtOH, and one object was replaced with a second, completely different object. Subject mice were returned to the arena and again allowed to explore both objects. The arena and all interaction objects were thoroughly cleaned with 70% EtOH between each subject mouse and before the first trial of the day. Novelty preference index was calculated on the second trial, defined as percent of time spent interacting with the novel object divided by total time spent interacting with either object.

### Sucrose preference test

Sucrose preference test was performed as previously described (PMID: 28462942). Following the novel object recognition test, mice were single housed with *ad libitum* access to food and two bottles: one with water and one with 1% w/v sucrose solution. For four days, liquid intake was measured daily by weighing the bottles, and the left/right positions of the bottles were swapped to avoid position preferences. Percent sucrose preference is expressed as (Δweightsucrose)/(Δweightsucrose + Δweightwater) × 100.

### Tissue preparation and RNA sequencing

Mice virally manipulated with HSV-TRIM28^NFD^, -TRIM28^WT^, -TRIM28^VPR^, or HSV-GFP were used in RNAseq analysis. Mice were cervically dislocated and decapitated without anesthesia, and the brains were removed and sectioned into 1 mm coronal slices using brain matrices. Central tissue punch containing bilateral PFC (12 gauge; internal diameter, 2.16 mm) were snap frozen on dry ice and stored at -80 °C, as is routinely performed by our group^22,73,74^. RNA was extracted and purified using RNeasy (Qiagen, Hilden, Germany), and total RNA was quantified with the Qubit RNA HS Assay Kit (Thermo Fisher Scientific, Waltham, MA). RNA quality control assays were performed on the TapeStation 4200 (Agilent, Santa Clara, CA), and the average RNA integrity number for all samples exceeded 8.6. Ribosomal RNA depletion and library preparation (Illumina Ribo-Zero) was performed, and RNA-seq was carried out at Genewiz with the following configuration: 2 × 150 paired-end reads on an Illumina (San Diego, CA) sequencing platform (HiSeq 2500) with a sequencing depth of ∼52 million reads per sample (mean = 52 ± 0.45 million). Other overall sample sequencing statistics include the mean quality score (37.77 ± 0.05) and the percent of bases ≥ 30 (92.20 ± 0.19).

### RNA-seq quality control and gene quantification

Raw RNA-seq FASTQ files were subjected to quality control using FastQC (version 0.11.9) to assess read quality. Adapter sequences were removed using Trimmomatic (version 0.39) with further trimming off low-quality bases (16 bases and 3 bases from the head and tail, respectively). We also dropped the 4-base window of average sequencing quality < 15 and finally the reads less than 50 bases. After the quality control, high-quality reads were aligned to the Mus musculus GRCm39 reference genome using STAR (version 2.7.11a) with the recommended parameters. We employed TEtranscripts^39^ (version 1.09) in the quantification of both gene and TE expression levels by integrating genomic and RepeatMasker annotations (Hammell lab). After extraction of gene hit counts, the gene hit counts table was used for downstream differential expression analysis.

The RNA-seq count data were normalized, and dispersion estimates were obtained according to DESeq2’s standard pipeline. DEG lists were generated relative to the HSV-GFP virus condition, and all viral conditions were analyzed both separated by sex and with pooling across sexes.

Wald tests were performed to determine differentially expressed genes (DEGs) and differentially expressed transposable elements (DETEs). Gene and TE IDs with p-values < 0.05 were considered DEGs and DETEs respectively. Volcano plots with these DEGs and DETEs were assembled using ggplot in Rstudio. Raw and processed RNAseq gene expression data are available via the Gene Expression Omnibus data (accession number will be made available upon publication).

### Genomic annotation of TE and potential regulated genes

We annotated genomic features overlapping with the origins of DETEs in generated from the above analysis. The genomic features included in this analysis were the promoter, enhancer, histone H3K4 tri-methylation and CTCF binding site revealed by ENCODE project, as well as the exon, intron, UTR from GRCm39 gene annotations. For the overlapping enhancers, we obtained their potential regulated genes predicted using ChIP-seq data from EnhancerAtlas v2.0^47^. A complete list of all TE annotations for the HSV-GFP vs. HSV-TRIM28^WT^, HSV-GFP vs. HSV-TRIM28^NFD^, and HSV-GFP vs. HSV-TRIM28^VPR^ comparisons are available upon request to the authors.

### DEG analysis approach

DEG tables from DESeq2 and annotated DETE data tables from the above analysis were imported into Rstudio for further analysis. DETEs were categorized by class and plotted using the ggplot package. Rank-rank hypergeometric overlap testing was completed on unfiltered DEG tables with DETEs excluded using the RRHO package. GSEA analysis was conducted using WebGestalt^75^ with a range in category size from 5-300 genes and ontology terms were calculated regardless of FDR. Results were plotted as a volcano plot using ggplot, and a significance criteria of FDR < 0.05 was applied. DETEs were filtered for unique genomic origins and sorted by their predicted annotated genomic features, then plotted using the ggplot package. We compiled all DEGs with a known TE association, then applied an unadjusted p < 0.05 cutoff from expression data from our DESeq tables. DEGs associated with DETEs were analyzed using over-representation analysis using WebGestalt^75^, and the gene ontology results were plotted using ggplot.

### Statistical analysis

The three-way ANOVA tests on social behavioral data were conducted using the rstatix package in R, and simple main effects were reported. Due to the lack of variance as a function of biological sex, all behavioral data was condensed to combine male and female data for further analysis. All data was otherwise analyzed in GraphPad Prism 10. In all figures, results were expressed as mean ± standard error (S.E.M.). *In vitro* validation data was analyzed by one-way ANOVA with Bonferroni correction (Fig. 1c, e). Behavioral data was analyzed as two-way

ANOVA with Bonferroni correction (Fig. 2b-c, e-g, j-k: Fig. 6a-b), one-way ANOVA with Bonferroni correction (Suppl. Fig. 1b-d, f, h: Fig. 2b-c) and Wilcoxon signed rank test (Fig. 2j-k). The distribution of DETEs by class was compared by Chi-square goodness of fit test (Fig. 3d). *P*-value < 0.05 was considered statistically significant. Experimental sample sizes were guided by power analyses and previously published experiments from our group and others. Animals were randomly assigned to viral treatment conditions. Experimenters were blind to treatment and experimental analysis was performed by automated Ethovision software.

## Acknowledgements

This work was supported by R00DA045795, P30DA033934, R01DA058958, and Blick Scholar funds to PJH, a V30 training award from the VCU MD-PhD Enrichment Fund to RKK, and F31MH133309 to NLT.

## Author Contributions

RKK, CS, and PJH conceived and conceptualized experimental design and data interpretation. RKK, CS, NLT, NC, GMS, and XC performed all experimentation. RKK and CS analyzed RNA sequencing data. RLN packaged herpes simplex viruses. RKK, CS, and PJH contributed to manuscript preparation. All authors reviewed and approved final manuscript draft.

## Notes

### Competing Interest Statement

The authors have declared no competing interest.

