## Supplemental Figures for "Derepression of transposable elements in mouse prefrontal cortex disrupts social behavior"

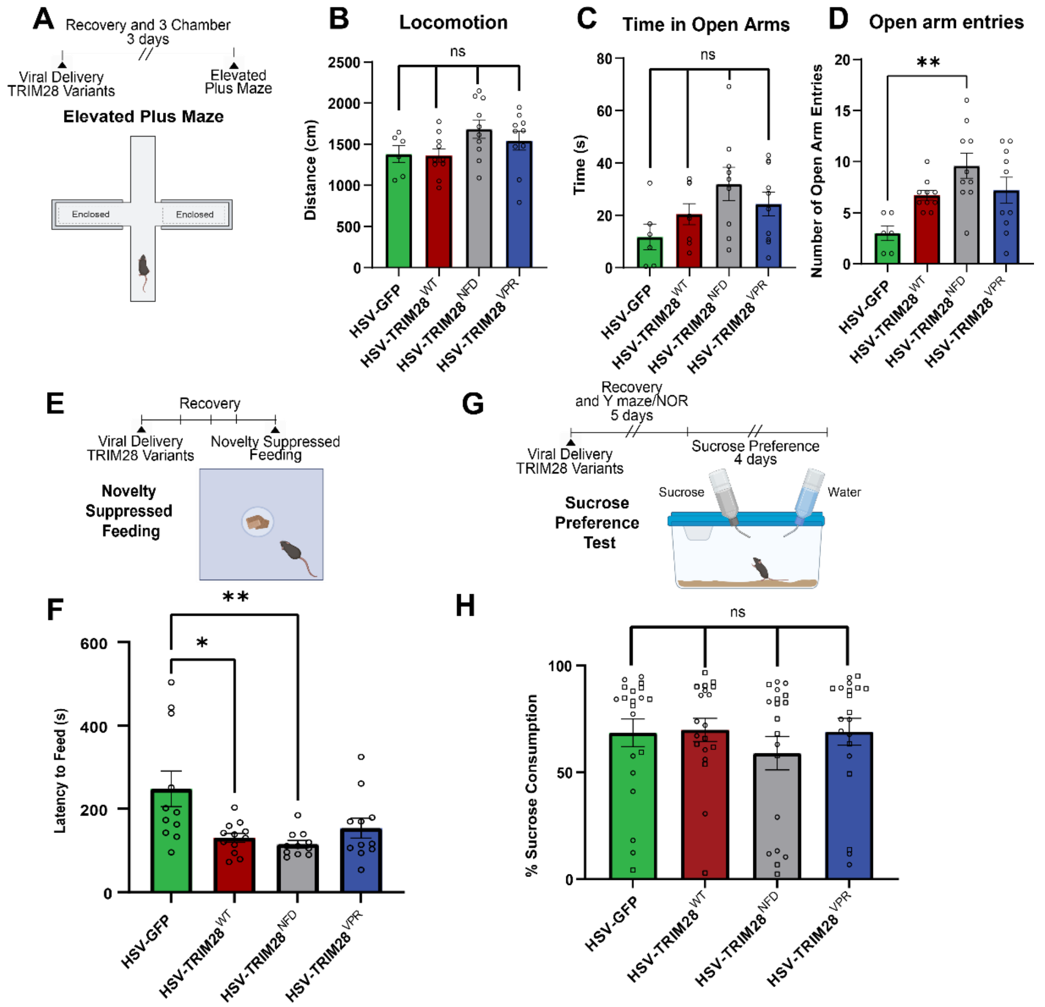


**Supplementary Figure 1: Disrupting PFC TRIM28 function does not impact anxiety-like behaviors.** a) Timeline and schematic of the elevated plus maze (EPM) (HSV-GFP: n = 6 males, HSV-TRIM28$^{WT}$: n = 10 males, HSV-TRIM28$^{NFD}$: n = 10 males, HSV-TRIM28$^{VPR}$: n = 10 males). b-d) Locomotion, time spent in the open arms, and entries into the open arms of the EPM is not significantly impacted by TRIM28 variants ($p > 0.05$, one-way ANOVA with Bonferroni correction). HSV-GFP-treated mice make significantly fewer entries into the open arms than HSV-TRIM$^{NFD}$-treated mice (panel d, $p = 0.0018$, one-way ANOVA with Bonferroni correction), but do not spend significantly less time in them (panel c, $p > 0.05$, one-way ANOVA with Bonferroni correction). e) Timeline and schematic of the novelty suppressed feeding arena (HSV-GFP: n = 11 males, HSV-TRIM28$^{WT}$: n = 12 males, HSV-TRIM28$^{NFD}$: n = 11 males, HSV-TRIM28$^{VPR}$: n = 11 males). f) HSV-TRIM28$^{WT}$ and -TRIM28$^{NFD}$-treated mice have a significantly lower latency to feed than HSV-GFP-treated mice ($p = 0.0104$ and $p = 0.0037$, respectively, one-way ANOVA with Bonferroni correction), but there is no significant difference between TRIM28 variant groups ($p > 0.05$ for all, one-way ANOVA with Bonferroni correction). g) Timeline and schematic of the sucrose preference test( HSV-GFP: n = 10 females, 10 males; HSV-TRIM28$^{WT}$: n = 10 females, 9 males; HSV-TRIM28$^{NFD}$: n = 10 females, 10 males; HSV-TRIM28$^{VPR}$: n = 10 females, 10 males) h) Viral treatment group does not significantly impact the percentage of sucrose consumed throughout the testing period ($p > 0.05$ for all, one-way ANOVA with Bonferroni correction). Squares indicate females; circles indicate males.

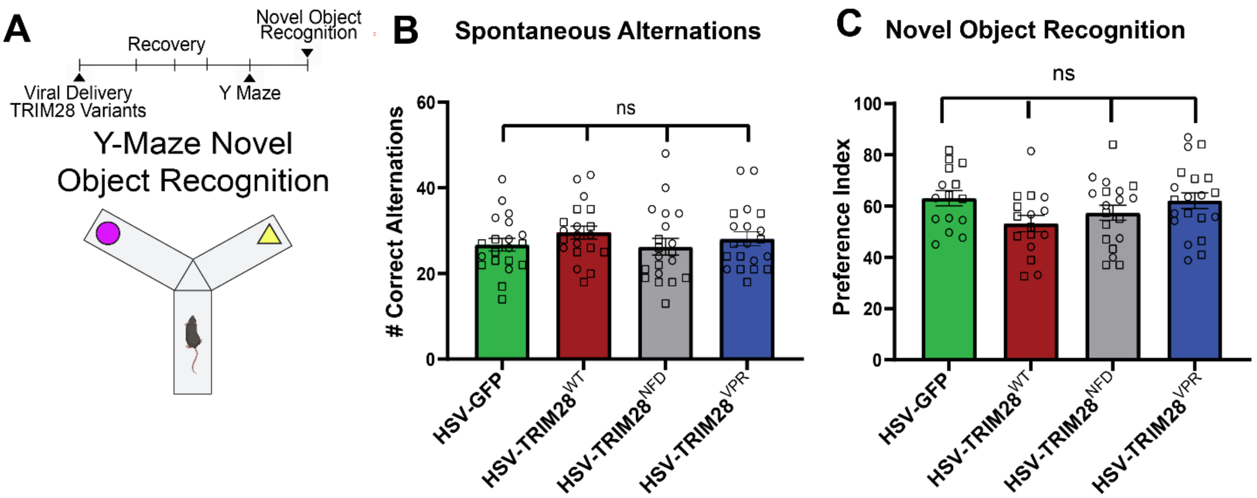


**Supplementary Figure 2: Disrupting PFC TRIM28 function does not affect non-social object recognition**. a) Timeline and schematic of novel object recognition in the Y-Maze (HSV-GFP: n = 10 females, 10 males; HSV-TRIM28$^{WT}$: n = 10 females, 9 males; HSV-TRIM28$^{NFD}$: n = 10 females, 10 males; HSV-TRIM28$^{VPR}$: n = 10 females, 10 males). b-c) Infusion of HSV-GFP or an HSV-TRIM28 variant to the prefrontal cortex (PFC) does not produce significantly different effects on the number of correct alternations or novelty recognition in the Y-Maze ($p > 0.05$ for all, one-way ANOVA with Bonferroni correction). Squares indicate females; circles indicate males.

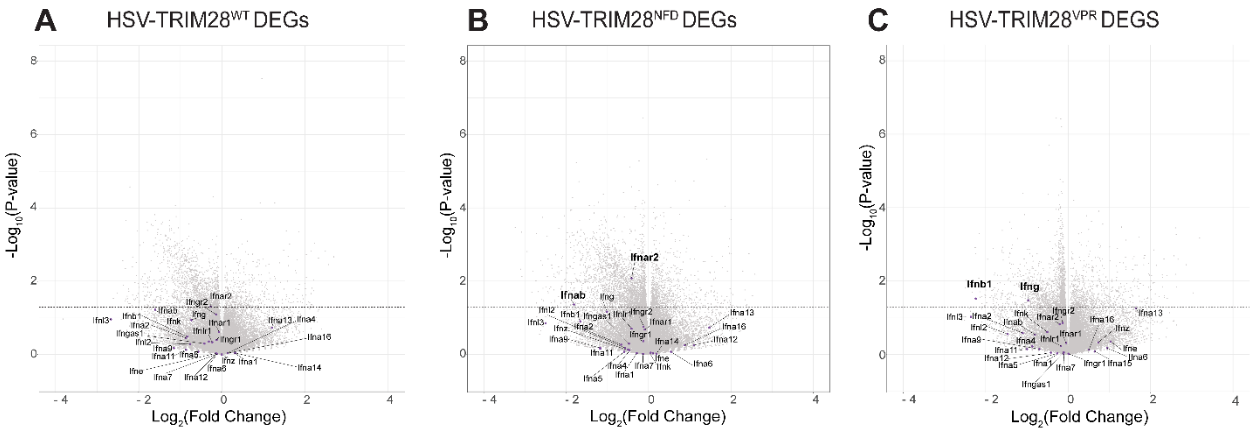


**Supplemental Figure 3: Variable downregulation of interferon cytokines by synthetic TRIM28 variants.** a-c) Volcano plots showing expression of interferon genes across HSV-TRIM28$^{WT}$, -TRIM28$^{NFD}$, and -TRIM28$^{VPR}$ conditions. No interferon cytokines are differentially expressed in HSV-TRIM28$^{WT}$. Only interferon alpha-B is downregulated in HSV-TRIM28$^{NFD}$. Both interferon beta and gamma are downregulated in HSV-TRIM28$^{VPR}$.

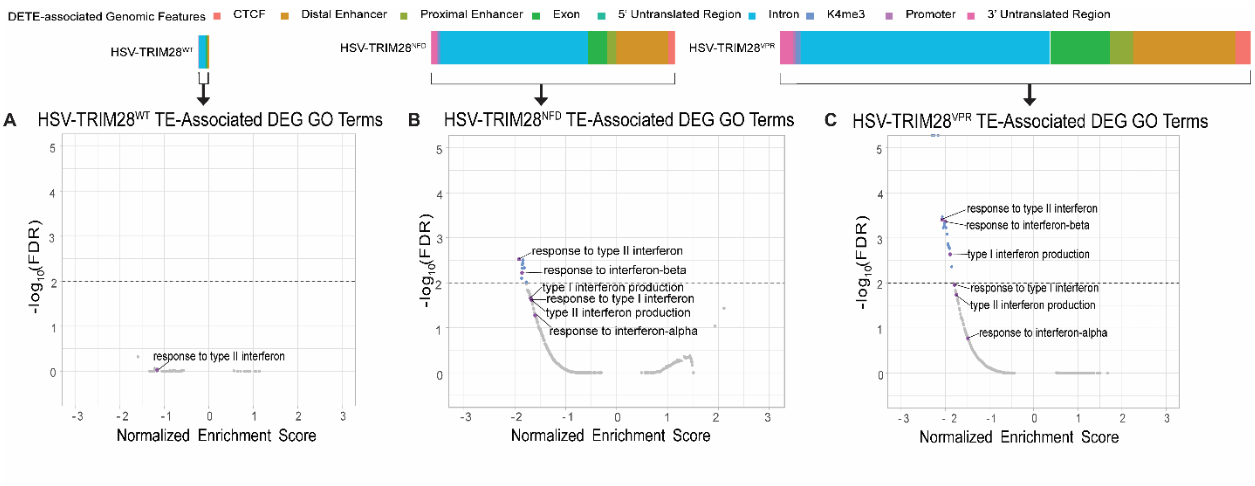


**Supplemental Figure 4: TE-associated DEGs are enriched for immune genes.** TE-associated DEGs were generated from searching DETEs through EnhancerAtlas and compiling a list of all significant DEGs associated with DETEs in each TRIM28 variant condition. a-c) Gene Set Enrichment Analysis was conducted on these TE-associated DEGs. Interferon related ontology terms are downregulated by HSV-TRIM28$^{VPR}$ to a greater extent than HSV-TRIM28$^{NFD}$. A significance criteria of FDR < 0.01 was applied.

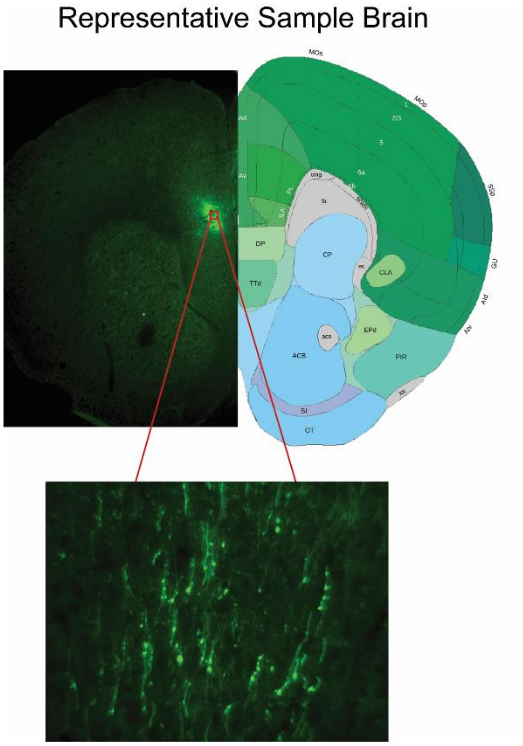


**Supplemental Figure 5: HSV transduction is not impaired by administration of exogenous interferon.** Co-delivery of HSV vectors with 10 ng each of interferon beta and gamma does not prevent viral transduction of PFC neurons. Representative PFC image shown.

| TE class | HSV-GFP vs HSV-TRIM28$^{VPR}$ | HSV-GFP vs HSV-TRIM28$^{NFD}$ | HSV-GFP vs HSV-TRIM28$^{WT}$ |
|---|---|---|---|
| Unclassified | 7 | 8 | 3 |
| Non-LTR | 166 | 67 | 8 |
| LTR | 229 | 127 | 36 |
| DNA | 108 | 57 | 14 |
| Total | 510 | 259 | 61 |

**Supplemental Table 4: Synthetic TRIM28 variants disrupt expression of all classes of transposable elements. HSV-**TRIM28$^{NFD}$ reflects derepression of TEs, while HSV-TRIM28$^{VPR}$ reflects active transcription of TEs.

| Genomic Feature | HSV-GFP vs HSV-TRIM28$^{VPR}$ | HSV-GFP vs HSV-TRIM28$^{NFD}$ | HSV-GFP vs HSV-TRIM28$^{WT}$ |
|---|---|---|---|
| CTCF | 9057 | 4006 | 80 |
| K4m3 | 3041 | 1562 | 35 |
| Distal Enhancer | 60594 | 30637 | 1016 |
| Proximal Enhancer | 13611 | 5359 | 142 |
| Exon | 35137 | 15950 | 692 |
| 5' Untranslated Region | 477 | 219 | 11 |
| Intron | 147483 | 87325 | 4393 |
| Promoter | 1410 | 579 | 16 |
| 3' Untranslated Region | 7681 | 3136 | 99 |
| Total | 278491 | 148773 | 6484 |

**Supplemental Table 5: Synthetic TRIM28 variants potentiate expression of transposable elements associated with distal enhancer and intronic genomic features.** The most abundant genomic origin associated with DETEs are distal enhancer and intronic regions.
